# Basal G-protein-gated inwardly rectifying potassium (Kir3/GirK) channels activity governs synaptic plasticity that supports dorsal hippocampus-dependent cognitive functions

**DOI:** 10.1101/2020.12.11.421735

**Authors:** Sara Temprano-Carazo, Souhail Djebari, Guillermo Iborra-Lázaro, Irene Sánchez-Rodríguez, Mauricio O. Nava-Mesa, Alejandro Múnera, Agnès Gruart, José M. Delgado-García, Lydia Jiménez-Díaz, Juan D. Navarro-López

**Affiliations:** University of Castilla-La Mancha, NeuroPhysiology & Behavior Laboratory, Centro Regional de Investigaciones Biomédicas, Facultad de Medicina de Ciudad Real, Spain; Neuroscience Research Group (NEUROS), University of Rosario, Bogotá, Colombia; Behavioral Neurophysiology Laboratory, Universidad Nacional de Colombia, Bogotá, Colombia; Pablo de Olavide University, Division of Neurosciences, Seville, Spain

**Keywords:** Kir3/GirK, hippocampus, neuronal excitability, LTP, LTD, metaplasticity, *ex vivo*, *in vivo*, Novel Object Recognition, Habituation test, Operant conditioning, mouse

## Abstract

G-protein-gated inwardly rectifying potassium (Kir3/GirK) channel is the effector of many G-protein-coupled receptors. Its dysfunction has been linked to the pathophysiology of Down syndrome, Alzheimer’s and Parkinson’s diseases, psychiatric disorders, epilepsy, drug addiction, or alcoholism. GirK channels are constitutively activated in the dorsal hippocampus contributing to resting membrane potential, and their synaptic activation compensates any excitation excess. Here, in order to elucidate the role of GirK channels activity in the maintenance of dorsal hippocampus-dependent cognitive functions, their involvement in controlling neuronal excitability at different levels of complexity was examined. For that purpose, basal GirK activity was pharmacologically modulated by two specific drugs: ML297, a GirK channel opener, and Tertiapin-Q, a GirK channel blocker. *Ex vivo,* using dorsal hippocampal slices, we studied pharmacological GirK modulation effect on synaptic plasticity processes induced in CA1 by Schaffer collateral stimulation. *In vivo*, we performed acute intracerebroventricular injections of both GirK modulators to study their contribution to CA3–CA1 synapse electrophysiological properties, synaptic plasticity, and learning and memory capabilities during hippocampal dependent tasks. We found that pharmacological disruption of basal GirK activity in dorsal hippocampus, causing either function gain or loss, induced learning and memory deficits by a mechanism involving neural excitability impairments and alterations in induction and maintenance of long-term synaptic plasticity processes. These results support the contention that an accurate control of GirK activity must take place in the hippocampus to sustain cognitive functions.

**Significance Statement:** The dorsal hippocampus mostly performs cognitive functions related to contextual/spatial associations. These functions rely on synaptic plasticity processes that are critically ruled by a finely tuned neural excitability. Being the downstream physiological effectors of a variety of G-coupled receptors, activation of G protein-gated inwardly rectifying K+ (GirK) channels induces neurons to hyperpolarize, contributing to neural excitability throughout the control of excitatory excess. Here, we demonstrate that modulation of basal GirK channels activity, causing either function gain or loss, transforms HFS-induced LTP into LTD, inducing deficits in dorsal hippocampus-dependent learning and memory. Together, our data show a crucial GirK activity-mediated mechanism that governs synaptic plasticity direction and modulates subsequent hippocampal-dependent cognitive functions.

## Introduction

G protein-gated inwardly rectifying potassium (Kir3, or GirK) channels are a family of K^+^ channels activated via ligand-stimulated G protein-coupled receptors (GPCRs) (Gonzalez et al., 2012). GPCR-GirK cascade can be started by many neurotransmitters inducing neurons to hyperpolarize, and contributing to control excitatory excess (Luscher and Slesinger, 2010). Being the downstream physiological effectors of a variety of receptors such as GABA (GABA_B_), serotonin (5HT-1A), adenosine (A_1_), muscarinic (M_2_), noradrenergic (α2), dopamine (D_2_, D_3_, and D_4_), opioid (μ, κ, and δ), cannabinoid (CB1) or somatostatin (Dascal and Kahanovitch, 2015; Glaaser and Slesinger, 2015), GirK channels have been pointed out as potential targets to be explored in many CNS disorders (Mayfield et al., 2015). Proof of that, GirK–dependent signaling disruption has been related to the etiology of several disorders such as Down’s syndrome, epilepsy, schizophrenia, autism, mood disorders, Alzheimer disease and drug abuse (Nava-Mesa et al., 2014; Slesinger et al., 2015; Tipps and Buck, 2015; Rifkin et al., 2017).

Genetic models have afforded knowledge about GirK channels functional role in normal and pathological conditions (Mayfield et al., 2015). However, these models raise deep concern about what compensatory mechanisms would occur due to global deletion of individual genes. In this sense, studies about the functional consequences of pharmacological modulation of GirK channels are scarce, since subunit-specific channel modulators development began until recently, when their crystal structure were resolved (Whorton and MacKinnon, 2013).

GirK channels consist of various combinations of four homologous subunits (GIRK1-4), although there is general agreement that GIRK1∕GIRK2 heteromultimers are the prototypical neural GirK channel (Lujan et al., 2014). GirK channels are specially relevant in dorsal hippocampus (Lujan and Aguado, 2015), a region that predominantly executes cognitive functions, since pharmacologic blockade or genetic deletion in this region interferes with memory acquisition and consolidation processes (Morris, 2007; Fanselow and Dong, 2010). GirK channels are constitutively active, contributing to resting conductances *ex vivo* (Chen and Johnston, 2005; Nava-Mesa et al., 2013) and gating long-term potentiation (LTP)(Malik and Johnston, 2017), a synaptic plasticity process considered the physiological substrate for memory formation in the hippocampus (Bliss et al., 2018). They have been also proposed to participate in memory consolidation processes by modulating hippocampal sharp waves and ripples (Trompoukis et al., 2020). Therefore, GirK channels are not only important for excitability regulation and normal synaptic transmission, but their basal activity might also modulate the predisposition of synapses to undergo subsequent neural plasticity processes (Sanchez-Rodriguez et al., 2017; Sanchez-Rodriguez et al., 2020), a phenomenon known as metaplasticity (Abraham, 2008). Changes in neural excitability may modify LTP induction threshold and even generate long term depression (LTD) (Cooper and Bear, 2012; Keck et al., 2017; Mayordomo-Cava et al., 2020; Sanchez-Rodriguez et al., 2020), a form of synaptic plasticity mainly associated to habituation forms of memory (Collingridge et al., 2010) and extinction of previous memories (Malleret et al., 2010). Even though basal GirK activity is a pivotal determinant for hippocampal principal neurons’ excitability (Drake et al., 1997; Chen and Johnston, 2005; Nava-Mesa et al., 2013; Malik and Johnston, 2017), its contribution to cognitive processes performed by dorsal hippocampus has not already been deeply investigated.

In this study, we pharmacologically modulated basal GirK activity in dorsal CA3–CA1 hippocampal synapse studying, in brain slices and behaving animals, excitability and synaptic plasticity processes and their upstream consequences on cognition. Interestingly, GirK activity enhancement-induced decreased excitability impaired LTP induction phase, while basal GirK activity reduction impaired LTP maintenance. As a result, deficits in habituation and recognition memories, as well as in operant learning, were observed. Since GirK channels are one of the main contributors to neural excitability control, they appear to be essential for normal hippocampal performance at synaptic level, playing a key role in LTP/LTD threshold regulation and in plasticity processes induction and maintenance supporting dorsal hippocampus-dependent cognitive functions.

## Methods

### Subjects

Experiments were performed on 105 C57BL/6 male adult mice (1–3 months old; 15-28 g (n = 30) and 24–32 g (n = 75) for *ex vivo* and *in vivo* experiments respectively). Animals were obtained from an official supplier (Charles River, France), housed in groups of 5 per cage and kept on a 12 h light/dark cycle with constant ambient temperature (21 ± 1°C) and humidity (50 ± 7%). Animals that underwent surgery were individually housed after such procedure. Food and water were available *ad libitum*. In all cases, animals were randomly allocated to experimental groups and experimenters were blind to treatment.

All experiments were performed in accordance with European Union guidelines (2010/63/EU) and Spanish regulations for the use of laboratory animals in chronic experiments (RD 53/2013 on the care of experimental animals: BOE 08/02/2013), and approved by local Ethics Committees of the Universities of Castilla-La Mancha and Pablo de Olavide. All efforts were made to minimize animal suffering.

### Ex vivo fEPSP Recordings: Hippocampal slice preparation

Hippocampal slices were prepared as described previously (Sanchez-Rodriguez et al., 2020) and coronal sections were selected as Schaffer collaterals are perfectly preserved (Xiong et al., 2017) (Fig. 1A). In brief, animals were deeply anesthetized with halothane (Fluothane, AstraZeneca, London, UK), briefly intracardially perfused with 1 ml oxygenated (95% O_2_ + 5% CO_2_) ice-cold (4-6°C) artificial cerebrospinal fluid (aCSF), with sucrose (234 mM; #84100; Sigma, Poole, UK) replacing NaCl to minimize damage, and decapitated. The brain was excised and rapidly immersed in oxygenated ice-cold aCSF containing (in mmol/L): 118 NaCl (#S9888; Sigma, Poole, UK), 3 KCL (#P3911; Sigma, Poole, UK), 1.5 CaCl2 (#499609; Sigma, Poole, UK), 1 MgCl2 (#208337; Sigma, Poole, UK), 25 NaHCO3 (#S6014; Sigma, Poole, UK), 30 Glucose (#G8270; Sigma, Poole, UK) and 1 NaH2PO4 (#S8282; Sigma, Poole, UK). Coronal brain slices (350-μm thick) containing the dorsal hippocampus were prepared with a vibratome (7000smz-2; Campden Instruments, Loughborough, UK) (Malik and Johnston, 2017). Slices were incubated, for at least 1 h, at room temperature (22°C) in oxygenated aCSF before recording.

**Figure 1.**
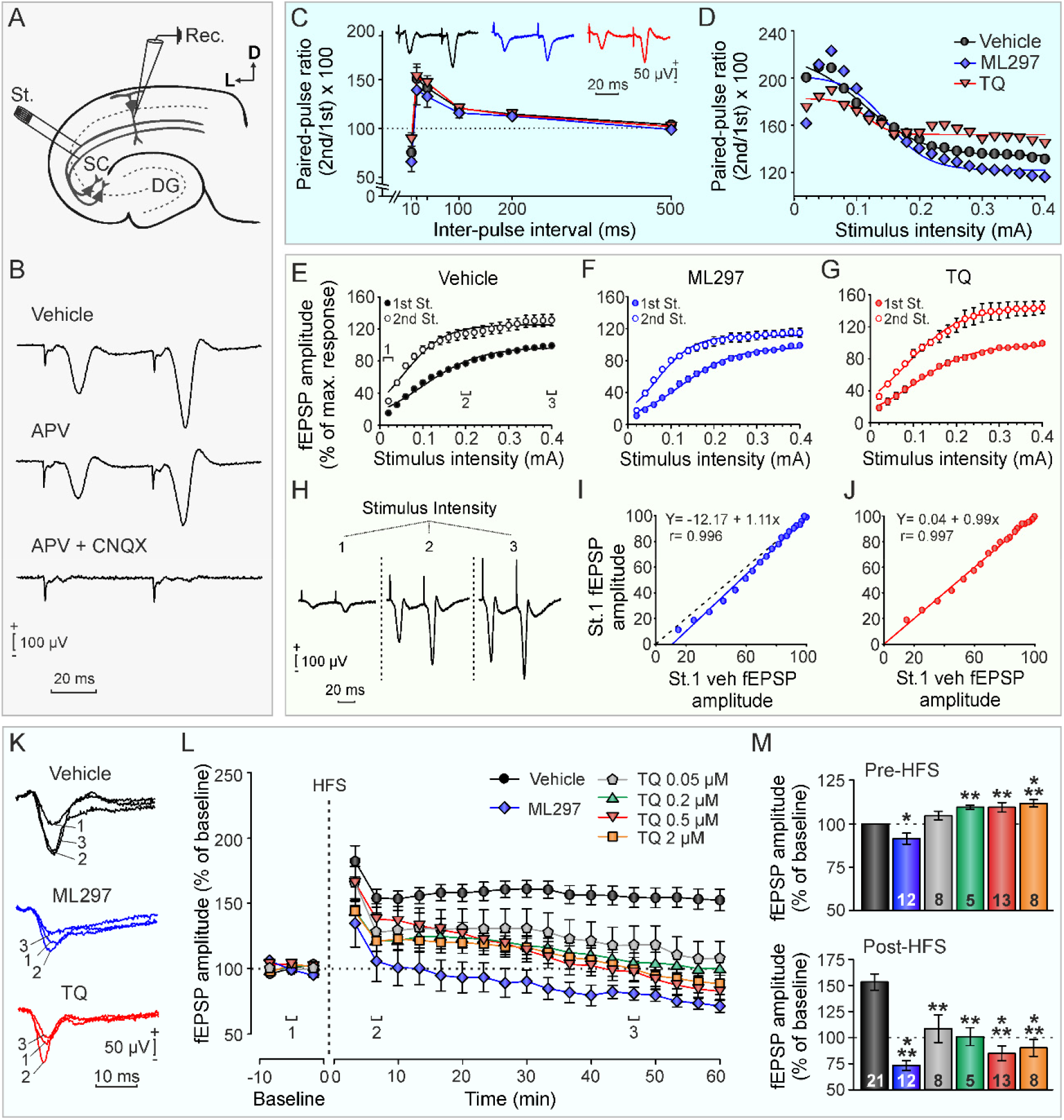
The modulation of basal GirK activity impairs LTP in the hippocampus. (**A**) Experimental design. The diagram illustrates the location of stimulation (St.) and recording (Rec.) electrodes in a hippocampal coronal slice. SC, Schaffer collaterals; DG, dentate gyrus. (**B**) Pharmacological characterization of the recorded fEPSPs in CA1. Perfusion of the slice with APV (50 μM; n = 10) did not affect the fEPSP, but it was completely abolished by CNQX (10 μM; n = 10), verifying that it was essentially mediated by glutamate-activated AMPA-kainate receptors. (**C**) Paired-pulse facilitation (PPF) curve at the CA1-CA3 synapse in slices perfused with aCSF alone (vehicle), ML297 (10 μM) and TQ (0.5μM). PPF was evoked by ~35% of the current amount required to evoke a saturating response. Averaged fEPSP paired traces for each pharmacological condition were collected at interstimulus intervals of 10-500 ms. Data are expressed as mean ± SEM amplitude of the second fEPSP expressed as the percentage of the first [(second/first) x 100] for each of the interstimulus used in this test (PP ratio) for each experimental group. (**D**) Evolution of the paired pulse ratio [(second/first) x 100] with increasing stimulus intensity from 0.02 to 0.4 mA. For the input/output (I/O) curves, Schaffer collaterals were also stimulated with paired pulses at intensities from 0.02 to 0.4 mA. (**E**), (**F**) and (**G**) relationship between the intensity of stimuli and the amplitude of the fEPSPs evoked in CA1. (**H**) Representative recordings are shown for 0.02 (1), 0.2 (2) and 0.4 (3) mA in control slices. **(I)** and **(J)**, fEPSP values evoked by the paired pulses in the different experimental groups vs control (x-axis, vehicle and y-axis, experimental group). (**K**) Representative averaged (n = 20) traces of fEPSPs recorded in the CA1 area by stimulation collected before HFS (1, baseline), 6 min after HFS (2), and about 46 min after HFS (3) in vehicle, ML297 (10 μM) and TQ (0.5μM)-treated slices. (**L**) Time course of LTP evoked in the CA1 area after an HFS session in controls (vehicle), ML297 (10 μM) and different TQ concentrations (0.05-2 μM). Bath application of ML297 completely prevented LTP induction. Perfusion of different concentrations of TQ also altered induction as well as maintenance of LTP. (**M**) fEPSP amplitude during baseline before HFS (upper bar plot), and the potentiation level in the last 10 min after HFS (bottom bar plot). The number of slices for each condition is indicated in the corresponding bar. Mean ± SEM is represented in L-M. Differences with respect to vehicle (control) were expressed as * *p* < 0.05, **; *p* < 0.01, ***; *p* < 0.001.

For recording, a single hippocampal slice was transferred to an interface recording chamber (BSC-HT and BSC-BU; Harvard Apparatus, Holliston, US) and perfused continuously with aCSF. Extracellular field potentials from de CA1 pyramidal neurons were recorded using a borosilicate glass micropipette (1–3 MΩ; RRID:SCR_008593; World Precision Instruments, Sarasota, US) filled with aCSF positioned on the slice surface in the *stratum radiatum* of CA1 and connected to the headstage of an extracellular recording amplifier (NeuroLog System; Digitimer, Hertfordshire, UK). The synaptic responses were evoked by paired pulses stimulation applied at 0.2 Hz on the Schaffer collateral pathway through a tungsten concentric bipolar stimulating electrode (World Precision Instruments, Sarasota, US) using a programmable stimulator (MASTER-9; A.M.P.I, Jerusalem, Israel) (Fig. 1A). Biphasic, 60-μs-long, square-wave pulses were adjusted to ~35% of the intensity necessary for evoking a maximum field excitatory postsynaptic potential (*f*EPSP) response. For paired-pulse facilitation (PPF) protocol, stimulus intensity was set to ~35% of the intensity for evoking a maximum *f*EPSP response was set to avoid population spikes; pairs of stimuli were then delivered at different interstimulus intervals (10, 20, 40, 100, 200, 500 ms). For input/output (I/O) curves, two stimuli of increasing intensity (0.02 mA-0.4 mA) were delivered at 40-ms interstimulus interval. For long term potentiation (LTP) induction, a high frequency stimulation (HFS) protocol was used and consisted of five 1-s-long 100 Hz trains delivered at 30-s inter-train interval. Baseline values of *f*EPSPs amplitude evoked at the CA3–CA1 synapse were collected at least 10 min prior to LTP induction. After LTP induction, *f*EPSPs were recorded during at least 60 min to evaluate early (E-LTP) and late LTP (L-LTP) phases (*i.e.*, induction and maintenance phases).

### In vivo experiments: Surgery

Experimental procedures used to record hippocampal *f*EPSP in freely behaving mice have been described in detail elsewhere (Gruart et al., 2006; Sanchez-Rodriguez et al., 2017) (Fig. 2A). Briefly, mice were anesthetized with 4-1.5% isoflurane (induction and maintenance, respectively; #13400264, ISOFLO®, Proyma S.L., Ciudad Real, Spain) delivered using a calibrated R580S vaporizer (RWD Life Science, Dover, US; flow rate: 0.5 L/min oxygen). Buprenorphine was administered intramuscularly as analgesic during and after surgery (0.01mg/kg; # 062009, BUPRENODALE®, Albet, Barcelona, Spain). Animals were implanted with bipolar stimulating electrodes aimed at the right Schaffer collateral-commissural pathway of the dorsal hippocampus (Fig. 2A; 2 mm lateral and 1.5 mm posterior to bregma; depth from brain surface, 1.0-1.5 mm (Paxinos and Franklin, 2001)), and with a recording electrode aimed at the ipsilateral *stratum radiatum* underneath CA1 area (1.2 mm lateral and 2.2 mm posterior to bregma; depth from brain surface, 1.0-1.5 mm). These electrodes were made from 50-μm, Teflon-coated tungsten wire (#W558415; Advent Research Materials, Eynsham, UK) and their precise location was verified histologically (Fig. 2A) and from *f*EPSPs profiles.

**Figure 2.**
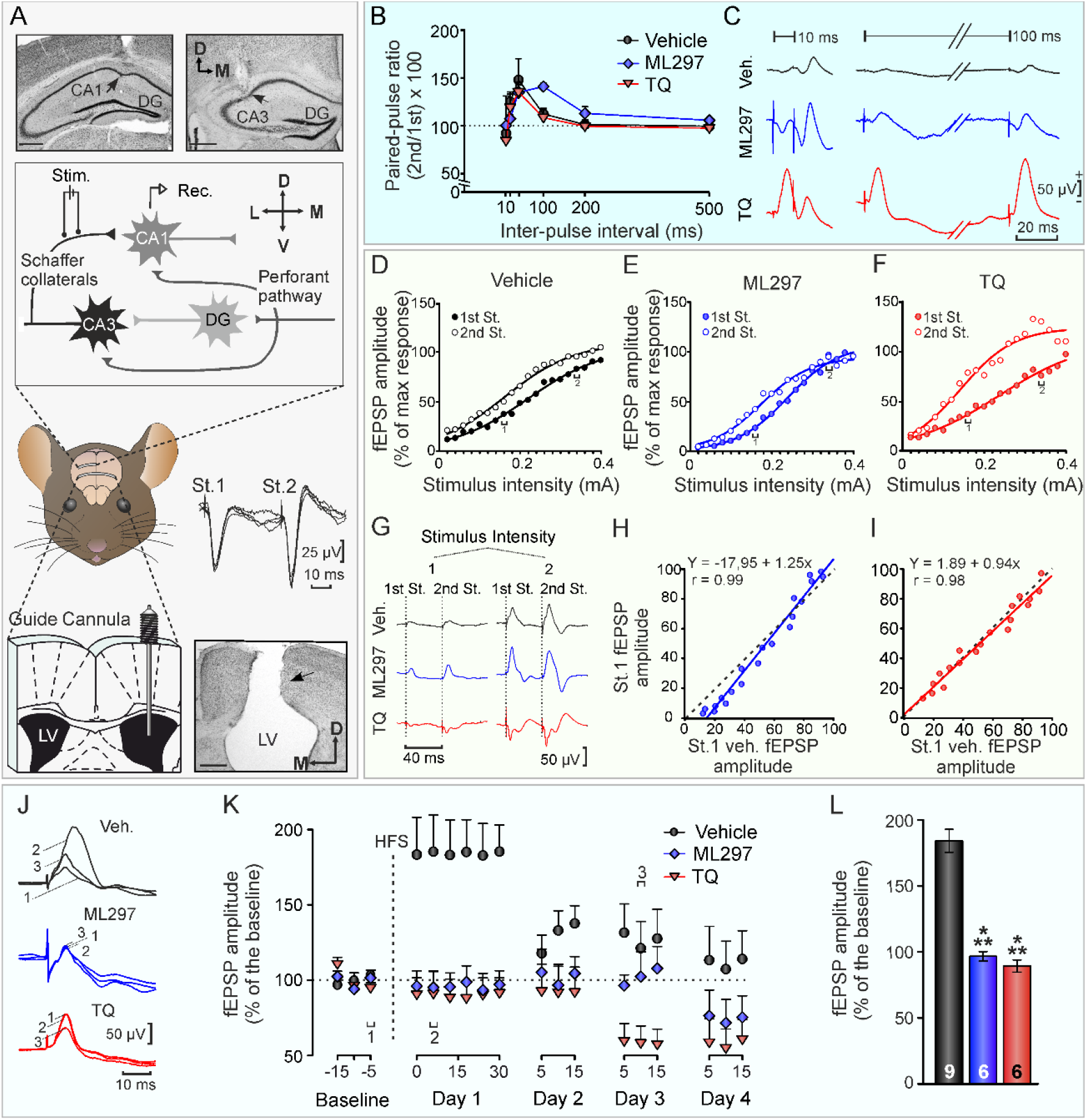
*In vivo* GirK activity modulation disrupted CA3-CA1 synaptic properties. (**A**) Mice preparation for in vivo chronic recording of fEPSPs evoked at the hippocampal CA3-CA1 synapse and *i.c.v.* drug administration. In the upper half, schema illustrates the location of the implanted recording (Rec.) and stimulating (Stim.) electrodes. Top photomicrographs show histological verification of electrodes position (indicated with black arrows). In the lower half, on the right, *f*EPSP profile evoked by paired pulses collected from a representative animal at intermediate stimulus intensities. Bottom left, a stainless steel guide cannula was implanted contralaterally to both electrodes, on the left ventricle. Bottom right photomicrograph serves as histological verification of cannula position (black arrow). Scale bars, 500 *μ*m. LV, Lateral Ventricle; DG, Dentate gyrus; PP, Perforant pathway; Sch., Schaffer Collateral; St., stimulus; D, dorsal; M, medial. (**B)** On the left, *in vivo* paired-pulse facilitation (PPF) curve at the CA1-CA3 synapse for *i.c.v.* injected mice with vehicle, ML297 and TQ. PPF was evoked by stimulating Schaffer collaterals at fixed current intensity (~35% of the current amount required to evoke a saturating response). Averaged (5 times) fEPSP paired traces for each experimental group were collected at interstimulus intervals of 10, 20, 40, 100, 200 and 500 ms. Data are expressed as mean ± SEM amplitude of the second fEPSP expressed as the percentage of the first [(second/first) x 100] for each of the 6 interstimulus intervals used in this test (PP ratio). (**C**) Representative examples (averaged ≥ 5 times) of fEPSPs evoked at the CA3-CA1 synapse at two different interstimulus intervals (10 and 100 ms) for each experimental group. **(D-F)** Input/output curves at the CA3–CA1 synapse for *i.c.v.* injected mice with vehicle, ML297 and TQ respectively were carried out with double pulses of increasing intensity at fixed inter-stimulus interval (40 ms). (**G**) Representative averaged (n = 5) records of *f*EPSPs recorded in the CA1 area following paired stimulation of the ipsilateral Schaffer collaterals at 2 different intensities (1, 0.16 mA and 2, 0.34 mA). **(H, I)** Scatter plots illustrating *f*EPSP values evoked by the first pulse in all experimental groups (x-axis, vehicle and y-axis, experimental group). The best linear fit is illustrated. **(J-L)** Evolution of *f*EPSPs evoked in the CA1 area by stimulation of Schaffer collaterals after an HFS session in freely moving mice. **(J**) Traces are representative examples of *f*EPSPs (averaged ≥ 5 times) collected from selected vehicle, ML297 or TQ *i.c.v.* injected mice evoked by pulses presented to the CA3‐CA1 synapse. *f*EPSPs were collected before (baseline) and after the HFS of Schaffer collaterals at the indicated times (see 1, 2, and 3 in the X axis of F2). (**K**) Graphs illustrate the time course of LTP evoked in the CA1 area by the paired pulses presented to Schaffer collaterals following HFS for vehicle (black circles), ML297 (blue diamonds) and TQ (red triangles) in for consecutive days. The HFS was presented after 15 min of baseline recording (dashed line). *f*EPSP amplitudes are given as the percentage of the baseline (100%) amplitude. Effects of HFS on *f*EPSPs evoked by the first pulses are illustrated. Note that only vehicle (control) mice presented a significant increase in *f*EPSP slopes following HFS when compared with baseline records. (**L**) Averaged *f*EPSPs of day 1 time points after HFS. The number of animals for each experimental group is indicated inside the corresponding bar. Mean ± SEM is represented in K-L. \*\*\**p* < 0.001.

For intracerebroventricular (*i.c.v.*) administration of drugs included in this study, animals were also implanted chronically with a blunted, stainless steel, 26-G guide cannula (Plastics One, Roanoke, US) aimed at lateral ventricle (0.5 mm posterior to bregma, 1.0 mm lateral to midline, and 1.8 mm below the brain surface) in the hemisphere contralateral to the one where stimulating and recording hippocampal electrodes were placed, as described elsewhere (Sanchez-Rodriguez et al., 2017) (Fig. 2A). Injections in freely moving mice were carried out with the help of a motorized Hamilton syringe at a 0.5-μL/min rate through a 33-G cannula, 0.5 mm longer than the implanted guide cannula and inserted inside it. Based on previous studies (Sanchez-Rodriguez et al., 2017; Sanchez-Rodriguez et al., 2019; Sanchez-Rodriguez et al., 2020), *i.c.v.* injections were done 24 h before *in vivo* electrophysiological protocols and 1 h before long term memory testing during behavioral learning tasks (see details below).

Finally, during surgical procedure, a bare silver wire (0.1 mm) was fixed to the skull as ground (Fig. 2A). Stimulating and recording electrodes and the ground were connected to a 6-pin socket that was then fixed to the skull with two small anchoring screws and dental cement. Mice were allowed a week for recovery before experimental sessions. Mice were routinely handled to minimize stress during experimental procedures.

### In vivo electrophysiological recordings in freely moving mice

To investigate the role of GirK channels in hippocampal functionality *in vivo*, we studied the excitability and functional capabilities of the CA3–CA1 synapse in alert behaving mice by generating I/O curves and testing synaptic plasticity through the induction of PPF and LTP by HFS of the hippocampal Schaffer collateral pathway, as previously described (Sanchez-Rodriguez et al., 2017)(Fig. 2). Only electrical recordings displaying clear field postsynaptic potentials (*f*PSP) components, without deterioration over time, and lacking signs of epileptiform activity (stimulus-evoked after-discharges, and/or ictal or post-ictal activity) were selected for analysis.

The *f*EPSPs were obtained from alert behaving mice using Grass P511 differential amplifiers through high-impedance probes (2×10^12^ Ω, 10 pF). Schaffer collaterals stimulation-evoked EPSPs were recorded from hippocampal CA1 area, while the animal was placed in a small (5×5×5 cm) box before (baseline (BL) values) and after *i.c.v.* injections. Electrical stimuli to Schaffer collaterals were 100-*μ*s long, square, biphasic pulses delivered either alone, paired, or in trains. As described for *ex vivo* experiments, for PPF protocol stimuli with intensity enough to evoke *f*EPSP with ~35% the maximum amplitude were delivered at 0-500 ms interstimulus intervals. For I/O curves, two stimuli with intensity ranging from 0.02 mA to 0.4 mA and were elicited at 40-ms interstimulus interval (Gruart et al., 2006; Sanchez-Rodriguez et al., 2017).

For LTP induction in behaving mice, stimuli intensity was also set at ~35% of that evoking maximal *f*EPSP amplitude. An additional criterion for selecting stimulus intensity for LTP induction was that a second stimulus, presented 40 ms after the first one (conditioning pulse), evoked a *f*EPSP with amplitude at least 150% the amplitude of the first one (Bliss and Gardner-Medwin, 1973). To obtain a BL, single 100-μs long, square, biphasic pulses delivered at 0.05 Hz to CA3–CA1 synapse were used to elicit *f*EPSPs and their amplitude were measured during 15 min before LTP. For LTP induction with HFS, six train clusters were delivered at 1-min intervals, each cluster consisted of five 100-Hz frequency, 100-ms long pulse trains delivered at 1-s intervals—therefore, a total of 300 pulses were used in a given LTP induction session (Gruart et al., 2006; Sanchez-Rodriguez et al., 2017). To avoid inducing large population spikes and/or epileptiform activity, stimulus intensity during HFS was the same as that used during BL. Immediately after LTP induction session, stimuli with the same parameters as for BL were delivered during 30 min. On successive days, BL stimulation parameters were used for 15-min long recording sessions. *f*EPSP amplitude data during HFS session and afterwards were normalized using BL *f*EPSP values collected on the first day as 100%; in this manner, it was possible to monitor *f*EPSP amplitude evolution during induction and maintenance phases of LTP (E-LTP and L-LTP).

### Behavioral experiments

#### Open Field Habituation task

Habituation to a novel environment in rodents is a type of non-associative hippocampal-dependent learning that can be measured as a change in exploration or in locomotor activity after re-exposure (Leussis and Bolivar, 2006). In the present work, to test this form of learning and its relation to hippocampal GirK signaling, mice were exposed twice to an open field (OF habituation task) as described elsewhere (Sanchez-Rodriguez et al., 2020). Briefly, animals performed two consecutive trials, one trial per day: training and habituation (or retention) trials on day 1 and 2 respectively (Fig. 3A). During training trial, mice were initially exposed to the OF. Upon 24 h later, habituation was tested (retention or habituation trial) by re-exposure of the animals to the same OF. *I.c.v.* injections were performed 1 h before retention testing. Drug concentrations used for *i.c.v.* injections had been previously shown not to affect motor activity in mice (Sanchez-Rodriguez et al., 2020). Exploratory behavior was tested using an actimeter (AC-5, Cibertec, Madrid, Spain), consisting of an infrared system to detect animal movements in a square white acrylic box (35×35cm×25 cm). On each trial, mice were placed at random in one of the four corners of the box and allowed to explore it for 15 min. Horizontal (X and Y axis crossing summation), vertical (Z axis crossing) and total (X, Y, Z axis crossing summation) movements were recorded. Recorded data were analyzed using MUX_XYZ16L software (Cibertec, Spain). The apparatus was cleaned with 70% ethanol to remove odors and allowed to dry completely before each animal was tested.

**Figure 3.**
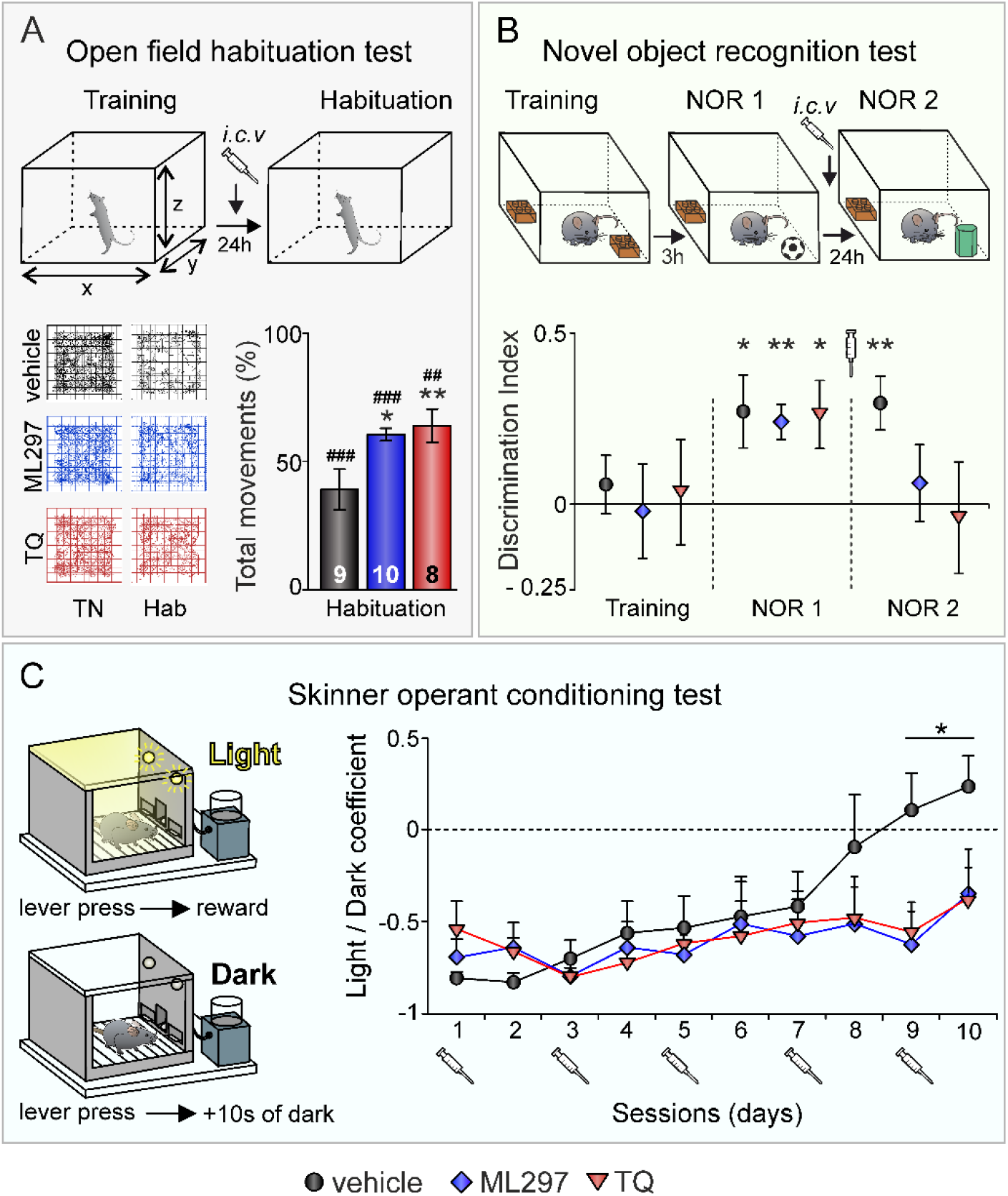
GirK modulation alter non-associative and associative learning. Experimental design for open field habituation (A), novel object recognition (B) and Skinner operant conditioning tasks (C) are represented in A-C. (**A**) Left bottom pannels show mice movement trackings during training (TN) and habituation (Hab) sessions for vehicle (control; in black), ML297 (in blue), and TQ (in red). Histogram represents total activity of each experimental group during habituation session expressed as percentage of exploration during training (100%). The number of animals for each experimental group is indicated in the corresponding bar. (**B**) Novel Object Recognition (NOR) test consisted of one training and two NOR testing sessions. During training two identical objects (yellow Lego pieces) were placed in the arena and animals explored for 5 min. After 3 h one object was replaced by a new one (mini-football ball) for the first test session (NOR1). After 24 h a second test (NOR2) was conducted with the familiar object and a new novel one (green cylinder). Animals were *i.c.v.* injected with vehicle, ML297 or TQ 1 h before NOR2 testing. Discrimination index represents the difference between time spent exploring novel and familiar objects. **(C**) Mice were trained in a Skinner box to press a lever to obtain condensed milk as reward with a fixed-ratio (1:1) schedule. Animals were trained with two programs of increasing difficulty. First, they had to acquire a fixed-ratio (1:1) schedule until obtaining, as a criterion, 20 times in 2 successive sessions. Afterward, animals were transferred to a light/dark paradigm in which lever presses were reinforced only when a light bulb was switched on. Lever presses performed during the dark period delayed the start of the lighted period for 10s. Light/dark coefficient (y-axis) was calculated as follows: (number of lever presses during the lighted period − number of lever presses during the dark period)/total number of lever presses. *I.c.v.* injections were carried out in odd sessions. A-C. Data represent mean ± SEM., \**p* < 0.05, \*\**p* < 0.01 (differences *vs.* vehicle (control) habituation), ## *p* < 0.01, ### *p* < 0.001 (differences *vs.* Training).

#### Novel Object Recognition task

Formation of object recognition (OR) memory is totally dependent on CA3–CA1 synaptic functionality in behaving mice (Clarke et al., 2010). Therefore, to further study the role of GirK channels on hippocampal-dependent learning, a Novel OR (NOR) task was conducted to test recognition memory. The task is based on the preference of mice for exploring a novel object over a familiar one, indicating then memory of the familiar object previously explored. Here, NOR was tested in a uniformly illuminated open-field arena (30×40×40 cm) and performed in 3 consecutive days (Fig. 3B) as previously described (Sanchez-Rodriguez et al., 2017). Briefly, on day 1 three habituation trials to the empty arena were performed (5-min trials, separated by 1.5 h resting periods). On day 2, two identical objects were placed in the center of the arena and animals were allowed to explore for 10 min (acquisition/training session). Both 3 and 24 h later, one object was replaced by a new one to perform two 10-minute test sessions (NOR1 and NOR2 sessions, see details below) to test short term memory (STM) and long term memory (LTM) respectively (Clarke et al., 2010; Sanchez-Rodriguez et al., 2017). New objects were different in shape and color but made of the same material (plastic) and with similar general dimensions. New objects and positioning of new objects were counterbalanced through all experiments to avoid bias. For assessment of STM retention and to determine if subjects were able to learn the NOR task and to properly consolidate and retrieve OR memory, exploratory behavior toward familiar and novel objects was quantified (NOR1 test). Only data from animals that performed successfully (≈94% of subjects) during NOR1—*i.e.*, explored the new object for longer than the familiar one—were analyzed and included in NOR2 test. To test LTM retention 24h after training session the novel object used at NOR1 was again replaced by a new novel one. Exploratory behavior towards novel and familiar objects was again measured. To analyze the impact of GirK channel activity modulation on LTM retention, *i.c.v.* injections of either vehicle, TQ, or ML297 were performed 1 h before the NOR2 test as described above. Learning was determined by quantification of the relative exploration time of each object and calculation of a Discrimination Index (DI), defined as the difference in exploration time between the two objects, O1 and O2 (TO1 and TO2, respectively), divided by the total time spent exploring both objects: (TO1-TO2)/(TO1+TO2). If we consider O1 to be the Novel Object, a DI greater than 0 will indicate exploration preference for such object.

Sessions were video recorded using a camera placed above the arena. Object exploration was scored by an experienced observer considering exploration only when the subject directed its nose to a given object at a distance ≤ 1 cm and/or touched/sniffed such object. When the subject used the object to prop itself up to explore the environment or accidently touched the object while heading in another direction the behavior was not computed as object exploration. To avoid the presence of olfactory trails, objects and arena were always thoroughly cleaned with ethanol 70% solution in between trials.

#### Operant conditioning task

Operant conditioning is a type of instrumental associative learning involving many cortical circuits activity, including the hippocampus (Jurado-Parras et al., 2016). In the present work we checked whether GirK activity modulation could affect acquisition of an associative learning task. Operant conditioning was conducted in two steps (Fig. 3C): a training phase and a light/dark test as previously described (Hasan et al., 2013). Both training and testing took place in a Skinner box (20.5×19.5×22.5 cm) located inside a sound-attenuating chamber (46×71.5×43 cm; Campden Instruments, Loughborough, UK). The operant chamber was equipped with a house light, two levers on the opposite wall with two lights on top of them, a feeder module between both levers and a liquid dispenser coupled to a pump that could deliver 10% condensed milk by lever pressing. Before training, mice were handled daily for 7 days and food-deprived to obtain a weight 85-90% of their free feeding weight. 20 min training sessions were done in successive days. Animals were trained to lever pressing using a fixed-ratio (FR 1:1) schedule to receive 20 μL-drops of condensed milk from the liquid dispenser. Turning house light on and off signalled each session start and finish, respectively. Mice were maintained on this 1:1 schedule until they reached criterion, pressing the lever ≥ 20 times/session in two successive sessions. Criterion was typically reached after 5-7 daily sessions.

Mice that reached training phase (fixed ratio 1:1) criterion were further conditioned using a light/dark (L/D) protocol. Animals were allowed a maximum of ten days to reach criterion. In L/D test animals were placed in the box for a maximum of 20 min in alternate periods of light and darkness of 20 s each. Only lever presses performed during the light period were reinforced with the liquid reward (success). In contrast, lever presses during the dark period were not rewarded (failure) and extended the period in 10 additional seconds. As this complex operant conditioning has been shown to rely on additional cortical regions than hippocampus (Fernandez-Lamo et al., 2018) and learning needs many sessions to be acquired, we decided to administrate the drugs sparingly along the test. *I.c.v.* injections of vehicle, TQ (2 μM), or ML297 (1.5 mM) were performed on alternating days of the L/D protocol (starting on session 1) 1 h before the session began. The L/D protocol lasted 10 days including one session/day/animal and the L/D coefficient [L/D coefficient = (number of lever presses during the light period – number of lever presses during the dark period) / total number of lever presses)] was calculated to determine the learning rate for each animal. In this case, the animal had to press the lever at least an equal number of times during the light (successes) and the dark (failures) periods; L/D ≥ 0) for 2 successive sessions to complete the task (criterion). Mice did not exhibit any motivational changes or motor deficits that could influence performance prior to the training phase or due to repeated *i.c.v.* injections of either ML297 or TQ during L/D testing. To avoid the presence of any odors the equipment was cleaned with 70% ethanol after each animal performance. Conditioning programs, lever presses, and delivered reinforcements were controlled and recorded by a computer, using ABET II Software (Campden Instruments, Loughborough, UK).

#### Histology and immunohistochemistry

As indicated above, the prototypical neural GirK channels in the hippocampus are GIRK1∕GIRK2 heteromultimers (Lujan et al., 2014). To study the level of expression of both GIRK subunits in the dorsal hippocampus, at the end of operant conditioning experiments in the Skinner box, mice were deeply anesthetized with ketamine/xylazine administered intraperitoneally (75/10 mg/Kg; KETALAR®, Pfizer, Spain and ROMPUM®, Bayer, Spain) and perfused transcardially with 0.9 % saline followed by 4 % paraformaldehyde in phosphate-buffered saline (PBS, 0.1 M, pH 7.4). This histological protocol also allowed to verify the proper location of implanted electrodes and cannulas. Mice brains were removed and cryoprotected with 30% sucrose in PB. Coronal sections (40 *μ*m) were obtained with a sliding freezing microtome (Microm HM 450, Walldorf, Germany) and stored at −20°C in 50 % glycerol and 50 % glycerol 50 % PBS until used. Sections including implantation sites in the hippocampus were mounted on gelatinized glass slides and stained using the Nissl technique with 0.25% Thionine to determine the location of stimulating and recording electrodes and/or the implanted cannula (Fig. 2A).

For fluorescence immunohistochemistry, free-floating sections were treated for 45 min with 10 % normal donkey serum (NDS; RRID:AB_2810235, Sigma Aldrich, Poole, UK) in Tris-buffered saline (TBS) containing 0.1 % Triton X-100 (TBS-T; #T8532, Sigma, Poole, UK), and subsequently incubated overnight at room temperature with polyclonal rabbit anti-GIRK1 (1:400; RRID:AB_2571710, Frontier Institute, Hokkaido, Japan) or polyclonal rabbit anti-GIRK2 (1:500; AB_2040115, Alomone Labs, Jerusalem, Israel) primary antibodies prepared in TBS-T with 0.05 % sodium azide (#S/2360/48, Fisher Scientific, New Hampshire, US) and 5% NDS. The following day, sections were washed with TBS-T (3 × 10 min) and then incubated for 2 h at room temperature with 1:150 dilutions of FITC-conjugated donkey anti-rabbit (RRID:AB_2315776, Jackson Immuno Research, West Grove, US) in TBS-T. After several washes with TBS (3 × 10 min), the tissue was incubated for 5 min in 0.01 % DAPI (#sc-3598, Santa Cruz Biotechnology, Santa Cruz, US) in TBS. Finally, sections were washed with TBS (3 × 10 min), mounted on gelatinized glass slides, dehydrated and coverslipped using a fluorescence mounting medium (#S3023, Dako mounting medium, Agilent, Santa Clara, US).

#### Image analysis

Images were acquired by confocal microscopy at 10X magnification using a laser scanning microscope (LSM 800, Carl Zeiss, Jena, Germany). GIRK1 and GIRK2 subunit expression were analyzed with ImageJ software (RRID:SCR_003070, NIH, Maryland, US) by measuring the intensity of immunostaining (*i.e.*, optical density) in randomly-selected squares of approximately 15 × 15 μm through the *stratum lacunosum-moleculare* and the molecular layer of the dentate gyrus at the dorsal hippocampus, as described elsewhere (Sanchez-Rodriguez et al., 2020). GirK channels are mainly expressed in the dendrites of pyramidal neurons in the dorsal hippocampus preferentially in distal dendrites. Thus, immunolabeling for both GIRK1 and GIRK2 subunits is more noticeable along the *stratum lacunosum-moleculare* and the molecular layer of the dentate gyrus (Fernandez-Alacid et al., 2011). As previously described, potential variations in optic density between experimental groups could be measured more accurately in these two areas, due to the greater range of measurement values. Mean background level was calculated from four different squares located at non-stained areas (*i.e.*, corpus callosum) and subtracted from the measurements. The values of optical density were normalized with respect to the control (vehicle) group values.

#### Drugs

All chemicals used in this study were purchased from Abcam (Cambridge, UK) and dissolved in PBS with the help of a shaker and/or sonicator. For *ex vivo* experiments, drugs ML297 (a selective opener of GIRK1–containing channels; 10 μM; #ab143564) or the channel blocker Tertiapin-Q (TQ; 0.02–0.5 μM; #ab120432) were dissolved in aCSF and applied by superfusion to the slices at a rate of 3mL/min (Sanchez-Rodriguez et al., 2020). For *i.c.v.* injection, drugs were dissolved in 3 μL of vehicle to get a solution of ML297 (1.5 mM), (Sanchez-Rodriguez et al., 2017), or the blocker TQ (1.5 mM) (Yow et al., 2011; Sanchez-Rodriguez et al., 2019), and injected through the guide cannula at a rate of 0.5 μL/min as described above.

#### Data collection and analysis

*In vitro* and *in vivo* recordings were stored digitally on a computer using an analog/digital converter (CED 1401 Plus, Cambridge, UK). Data were analyzed with the Spike 2 program (CED, Cambridge, UK). As synaptic responses were not contaminated by population spikes, the amplitude (i.e, the peak-to-peak value in mV during the rise-time period) of successively evoked fPSPs was computed and stored for later analysis.

All calculations were performed using SPSS version 20 software (RRID:SCR_002865; IBM, Armonk, US). When the distribution of the variables was normal, acquired data were analyzed with two-tailed Student’s *t* test or one-way or two-way ANOVA with *time* and *treatment* as within- and between-subjects factors respectively, except in I/O experiments in which intensity was the repeated measure), and with a contrast analysis for a further study of significant differences. For repeated measures two-way ANOVA, Greenhouse Geiser correction was used and indicated in the text when sphericity was not assumed. Statistical significance was set at *p* < 0.05.

Unless otherwise indicated, data are represented as the mean ± SEM. Computed results were processed for graphical purposes using the SigmaPlot 11.0 package (RRID:SCR_003210; Systat Software Inc., CA, US). Final figures were prepared using CorelDraw X8 Software (RRID:SCR_014235; Ottawa, Canada).

## Results

GirK channels are constitutively activated in the dorsal hippocampus (Chen and Johnston, 2005) suggesting a critical role for cognitive processes that depends on such region. Hence the goal of this work was to clarify the contribution of basal GirK activity to the related cognitive learning processes mainly focused on hippocampal–dependent learning and memory and underlying long-lasting synaptic plasticity phenomena at CA1–CA3 synapse.

### Dorsal hippocampus needs for GirK basal activity to induce and maintain LTP

We started our study in a classical slice preparation including dorsal hippocampus (Fig. 1A), by testing the effect of GirK activity modulation on the main functional properties of CA3–CA1 synapse, PPF, I/O and LTP. Firstly, single pulse stimulation applied at Schaffer collaterals was adjusted to yield a large *f*EPSP with a latency of 3.5– 4 ms in the CA1 *stratum radiatum* free of population spikes (Fig. 1B). Perfusion of the slice with APV (50 μM; n = 10) did not significantly affect the fEPSP, but it was completely abolished by CNQX (20 μM; n = 10), verifying that it was essentially mediated by glutamate-activated AMPA-kainate receptors (Fig. 1B).

Next, we checked whether a typical short-term plasticity phenomenon at the Schaffer collaterals-CA1 pyramidal neurons, PPF (Zucker and Regehr, 2002), was altered by basal GirK activity modulation. This facilitation protocol also allowed to assess the presynaptic functionality and, therefore, changes in neurotransmitter release. We tested slices for enhancement of synaptic transmission evoked by PPF using a wide range of interstimulus intervals (from 10 ms to 500 ms) at a fixed intensity (~35% of the amount needed for evoking a maximum fEPSP response). As illustrated in Fig. 1C, modulation of GirK conductance induced no significant differences at any of the selected (10, 20, 40, 100, 200, and 500 ms) intervals (F_(3.7,38.7)_ = 0.42, *p* = 0.78, veh. n = 12; ML297, n = 6; TQ, n = 6). In addition, we examined the evolution of the paired-pulse ratio (PPR) at 40 ms interval, with increasing intensities, from 0.02 to 0.4 mA (Fig. 1D). No significant differences were found neither in the global evolution of PPR at different intensities (F_(38, 342)_ = 0.708, *p* = 0.903, veh. n = 11; ML297, n = 5; TQ, n = 5), nor in PPR for each intensity (*p* > 0.05 for all cases). Paired-pulse analysis indicate that short-term plasticity processes and presynaptic vesicle release probability are not affected by GirK basal activity.

The excitability of the CA3–CA1 synapse was then studied by I/O protocols. The stimulus intensity was increased in steps of 0.02 mA from 0.02 to 0.4 mA, maintaining a fixed 40-ms interval between paired pulses. Increasing intensity of electrical stimulation delivered at Schaffer collaterals was accompanied by growing eater amplitude of fEPSPs for first (F_(19, 190)_ = 272.159, *p* < 0.001) and second F_(19, 190)_ = 11.336, *p* < 0.001) stimulus (Fig. 1E, H; n = 11). Slices perfused with ML297 showed a significant decrease of fEPSPs evoked by second pulse (Fig. 1F; n = 5; F_(19, 266)_ = 3.371, *p* < 0.001) whereas those perfused with TQ presented an increase for the response evoked by second stimulus (Fig. 1G; n = 5; F_(19, 266)_ = 4.318, *p* < 0.001). The scatter plots in Figs. 1I and 1J compare fEPSP amplitudes of the first pulse collected from slices treated with vehicle during the I/O study (on the x-axis) with the corresponding values for ML297 or TQ perfusion (y-axis). Data were shifted and presented a linear slope > 1 (*b* = 1.11) for ML297 according to a decreased excitability, while data for TQ fitted to value ≈ 1 (*b* = 0.99) suggesting that any tendency to increase synapse excitability could be damped.

As we found GirK channels to modulate CA3-CA1 synapse excitability, before LTP induction, basal *f*EPSP amplitude was monitored during the perfusion of specific GirK drugs (Fig. 1K, L, M). Amplitude of synaptic responses was decreased by ML297 (*t*(35) = 2.56, *p* = 0.015; Fig. 1L, M) whereas TQ significantly increased *f*EPSPs amplitude except for the lower concentration (0.05 μM, *t*(23) = −1.54, *p* = 0.14; 0.2 μM, *t*(14) = −4.13, *p* = 0.01; 0.5 μM, *t*(38) = −3.39, *p* = 0.002; 2 μM; *t*(23) = −5.36, *p* < 0.001; Fig. 1L, M). Then, after baseline was stabilized, we examined the effects of GirK modulation on synaptic plasticity. HFS applied at Schaffer collaterals induced a robust synaptic potentiation at CA3–CA1 synapses in control hippocampal slices which remained stable for, at least, one hour (156 ± 1.7%, n = 21; F_(1.1, 22)_ = 44.3 Greenhouse Geiser correction, *p* < 0.001 *vs.* baseline, Fig. 1L). However, the presence of the GirK1–containing channels opener ML297 (10 μM) blocked LTP induction (73 ± 2.6%, n = 12; F_(1.2, 13.5)_ = 32.3 Greenhouse Geiser correction, *p* < 0.001, Fig. 1K, L, M) even causing a LTD (Fig. 1L; *post hoc* vs. baseline *p* < 0.001). Interestingly, LTP was also induced, although with less amplitude than in controls, and maintained during 30 min after HFS at different TQ concentrations (Fig. 1K, L, M; 0.05 μM, n = 8, 130 ± 5%, F_(10, 70)_ = 3.99, *p* < 0.001; 0.2 μM, n = 5, 122 ± 1.8%, F_(10, 40)_ = 8.36, *p* < 0.001; 0.5 μM, n = 13, 127 ± 3.3%, F_(1.6, 18.8)_ = 7.28 Greenhouse Geiser correction, *p* = 0.007; 2 μM, n = 8, 120 ± 2.7%, F_(10, 70)_ = 5.39, *p* < 0.001). 30 min after induction none of these potentiations could be maintained (Fig. 1L, M) and disappeared about 50 min post-HFS (Fig. 1M; 10 last minutes post-HFS *vs.* baseline, F_(5, 61)_ = 16.08, p < 0.001; *poshoc* 0.05 μM, *p* = 0.002; 0.2 μM, *p* = 0.003; 0.5 μM, *p* < 0.001; 2 μM, *p* < 0.001) even reaching values of synaptic depression for 0.5 μM. Hence, our results suggest that an optimal range of GirK activity is required for LTP induction and maintenance, and channel malfunction creates an alteration that disturbs this critical form of plasticity in the dorsal hippocampus.

### In vivo LTP induction in the dorsal hippocampus requires of CA3–CA1 excitability regulation exerted by GirK channels

Long-term synaptic plasticity phenomena (LTP/LTD) that take place in the dorsal hippocampus has been shown to primarily perform cognitive functions (Fanselow and Dong, 2010). The induction threshold for LTP/LTD depends on hippocampal excitability level (Luscher and Malenka, 2012; Keck et al., 2017) controlled, among others, by GirK channels (Luscher and Slesinger, 2010) as we also found here *ex vivo.* Then, before studying the impact of channel modulation in related hippocampal-dependent learning and memory functions in mice, the next step was to investigate *in vivo* the role of hippocampal GirK activity on CA3–CA1 excitability and subsequently, in short– and long-lasting synaptic plasticity changes. To address this question, we took advantage of *in vivo* recording techniques and studied functional properties of the dorsal CA3–CA1 hippocampal synapse by I/O, PPF and LTP protocols in behaving mice (Fig. 2).

To study CA3–CA1 excitability, I/O protocol consisted of paired electrical stimuli with an interval of 40 ms and increasing intensity (range 0.02–0.4 mA in steps of 0.02 mA) was applied at Schaffer collaterals to evoke a large *f*EPSP in the CA1 pyramidal cells (Fig. 2A, G). *f*EPSPs from control animals showed the characteristic increasing sigmoid-like curve for the first (F_(19, 247)_ = 58.68, *p* < 0.001) and second (F_(19, 247)_ = 19.35, *p* < 0.001) stimulus (Fig. 2D,G). The evolution of the first and the second *f*EPSP evoked by the same pair of pulses at the same range of intensities (0.02– 0.4 mA) was not significantly different for mice injected with ML297 (Fig. 2E, G) or TQ (Fig. 2F, G). Excitability of the CA3–CA1 pathway was evaluated by plotting *f*EPSP values evoked by the first pulse in experimental groups (Y-axis) against control values (X-axis) (Gruart et al., 2012). As we also found *ex vivo*, for ML297 group, values were shifted downward, and the slope of the linear fits were > 1 (*b* = 1.25), in accordance with a decrease in excitability (Fig. 2H), while for TQ group slope was < 1 (*b* = 0.94) and values were shifted upward, suggesting a slight excitability increase (Gruart et al., 2012; Sanchez-Rodriguez et al., 2017)(Fig. 2I).

Next, functional capabilities of the CA3–CA1 synapse were studied by checking a typical short-term plasticity phenomenon of this synapse, the paired-pulse facilitation (PPF; Fig. 2B, C)). Mice were tested for enhancement of synaptic transmission evoked by PPF using a wide range of interstimulus intervals (from 10 ms to 500 ms) at a fixed intensity (≈35% of the amount needed for evoking a maximum fEPSP response). As illustrated in Fig. 2B, all groups presented a significant (F_(2.75, 77.06)_ = 5.30, Greenhouse Geisser correction, *p* = 0.003) increase of the response to the second pulse at short (40–100 ms) time intervals. No significant differences between groups were observed at any of the selected (10, 20, 40, 100, 200, and 500 ms) intervals (F_(2, 28)_ = 1.02, *p* = 0.37), thus suggesting not only a normal short-term hippocampal plasticity but also that the drugs used in the present work are preferentially acting at postsynaptic locations, as we found *ex vivo*.

As neural excitability level regulates the induction threshold for LTP/LTD (Keck et al., 2017) and we observed this transformation *ex vivo* when GirK conductance was modified, we asked whether modulation of the GirK–dependent signaling would impair this type of synaptic plasticity *in vivo*. HFS applied at CA3 Schaffer collaterals induced a significant enhanced *f*EPSP amplitude in the *stratum radiatum* of dorsal CA1 (184 ± 9% of baseline; n = 9; Fig. 2J, K, L) during the 30 min following HFS (F_(1.43, 11.42)_ = 12.08, *p* = 0.003). However, LTP could not be induced when basal GirK activity was pharmacologically enhanced or reduced (Fig. 2J, K. L)(ML297, F_(8, 40)_ = 0.3, *p* = 0.963, n = 6; TQ, F(8, 40) = 0.924, *p* = 0.508, n = 6). Interestingly, modulation induced a depression of the synaptic response that was noticeable in ML-injected (75 ± 8% of baseline) and TQ-injected (58 ± 6% of baseline) animals 72 and 48 h after HFS, respectively (Fig. 2K). These results suggest that constitutive activity of GirK channels is critical for the induction and maintenance of synaptic plasticity processes in the dorsal hippocampus that support learning and memory in behaving mice.

### GirK activity modulation impairs non-associative and recognition hippocampal-dependent learning

Our electrophysiological results *ex vivo* and *in vivo* strongly suggested that constitutive GirK activity was essential to control both excitability of the CA3-CA1 synapse and plastic mechanisms involved in memory storage. Many associative and non-associative learning and memory tasks have been shown to rely on CA3–CA1 dorsal hippocampus functionality (Gruart et al., 2006; Gruart and Delgado-Garcia, 2007; Clarke et al., 2010; Fernandez et al., 2017). So, we then wanted to assess the contribution of basal hippocampal GirK activity to such cognitive processes.

First, we challenged our experimental groups on the open field (OF) habituation test. Such non-associative hippocampal-dependent learning (Leussis and Bolivar, 2006) is based on the rodents tendency to decrease exploratory behavior in response to continued or repeated exposure to a novel environment (*i.e.* habituation) as an OF (Sanchez-Rodriguez et al., 2020). In the acquisition trial (training), experimental groups showed no significant differences in exploratory activity (Fig. 3A). In contrast, due to the habituation process, movements were significantly reduced on the retention day in all animals (vehicle (n = 9): *t*(8) = 7.22, *p* < 0.001; ML297 (n = 10): *t*(9) = 11.27; *p* < 0.001; TQ (n = 8): *t*(7) = 4.65; *p* = 0.002) (Fig. 3A). However, the decrease was more noticeable for vehicle than ML297 or TQ-injected animals (ML297 *vs*. vehicle: *p* = 0.014, TQ *vs*. vehicle: *p* = 0.008) suggesting that basal GirK activity disruption (either an increase or decrease) in the dorsal hippocampus impairs consolidation of habituation memory.

Additionally we wanted to investigated the role of GirK activity in long-term memory, so we choose the object recognition (OR) memory test as it has been shown to be totally reliant on dorsal hippocampal activity in behaving mice (Clarke et al., 2010). The task relies on the innate curiosity of rodents to explore a novel object more than a familiar one. During training (Fig. 3B), animals spent a similar amount of time exploring each object, with a discrimination index (DI) near to 0 (*p* > 0.05 in all experimental groups; Fig. 3B). Then, 3 h after training, the NOR1 session was performed and mice showed a strong preference for exploring the novel object (vehicle (n = 16): DI = 0.27± 0.11, *t*(15) = 2.54, *p* = 0.023; ML297 (n = 10): DI = 0.24 ± 0.05, *t*(9) = 4.7, *p* = 0.001; and TQ (n = 6): DI = 0.26 ± 0.1, *t*(5) = 2.6, *p* = 0.047). 24 h later *i.c.v.* drug injections were performed, and after 1 h, mice underwent a new testing session (NOR2). Vehicle-injected mice (n = 16) showed DI values similar to NOR1 showing again a clear discrimination between novel and familiar objects for these control animals (DI = 0.3 ± 0.08, *t*(15) = 3.76, *p* = 0.002). However, DI values for ML297- and TQ-injected mice were not significantly different from training values (ML297 (n = 8): DI = 0.06 ± 0.11, *t*(7) = 0.55, *p* = 0.602; and TQ (n = 6): DI = −0.04 ± 0.16, *t*(5) = −0.25, *p* = 0.82), indicating that these animals were not able to recognize the new objects (Fig. 3B).

In summary, these experiments demonstrate that dysregulation of basal GirK activity also disrupts long-term memory processes (non-associative and recognition memories) that depend on dorsal hippocampus performance.

### Role of GirK channels in associative learning: Operant conditioning

Finally, as we found basal GirK activity to be essential for non-associative hippocampal dependent memory, we also wanted to assess its involvement in associative learning. For such purpose, an operant conditioning test was carried out as the hippocampus is involved in both acquisition and storage of this type of learning (Jurado-Parras et al., 2016). Mice were first trained in a Skinner box to obtain condensed milk every time they pressed a lever in daily sessions of 20 min (fixed ratio of 1:1, Fig. 3C). The criterion of successfully completion of the task was 20 lever presses in 2 successive sessions. After 10 sessions, 87.5% of trained mice completed the task (Fig. 3C). Animals that successfully reached the above criterion were challenged to a more complex operant conditioning task designed as a light/dark (L/D) conditioning test with 10 sessions. Pressing the lever was rewarded only during periods of 20 s in which a light bulb located above the lever was switched on (Fig. 3C). Then, mice were *i.c.v.* injected with GirK selective drugs every two sessions (Fig. 3C). Statistical analysis of the learning rate (measured by the L/D coefficient) showed that only vehicle-injected mice (n = 6) significantly progressed along testing sessions (F_(9, 45)_ = 9.11, *p* < 0.001).

Moreover, from the 9^th^ session, animals injected with vehicle showed significant differences in the L/D coefficient (F_(2, 16)_ = 4.03, *p* = 0.04) with respect to both ML297- (n = 6; post hoc *vs.* vehicle, *p* = 0.03) and TQ-injected groups (n = 7; post hoc *vs.* vehicle, *p* = 0.03) (Fig. 3C), indicating learning success for control mice. Since animals injected with GirK modulators did not exhibit any motor or motivational impairment, our findings cannot be attributed to any specific difficulty to move around in the Skinner box or to any evident hyperactivity or motor inactivity (Jurado-Parras et al., 2016). Hence, these data show that altering constitutive GirK activity at CA3–CA1 synapses by pharmacological modulation of the channel also results in a significant deficit in the ability to learn an associative operant task.

### GirK protein expression in dorsal hippocampus after *in vivo* basal activity modulation

As GirK channels are formed by GIRK1/GIRK2 heteromultimers in the hippocampus (Fernandez-Alacid et al., 2011), we then asked whether modulation of basal GirK activity would alter the expression of both subunits at the protein level. We analyzed the optical density of GIRK1 and GIRK2 subunits in immunostained sections through the *stratum lacunosum-moleculare* (SLM) and dentate gyrus molecular layer (MDG) of the dorsal hippocampus as GirK channels are mainly expressed in distal dendrites of hippocampal pyramidal neurons and immunostaining is more noticeable in both areas (SLM and MDG). Our results show that *i.c.v.* injections of ML297 produced a significant decrease in GIRK1 and GIRK2 staining (Fig. 4A, B, D) in the SLM (GIRK1, n = 15 slices, 72 ± 3.3% of control vehicle values, *post hoc vs*. control, *p* < 0.001; GIRK2, n = 15 slices, 78 ± 4.8% of control vehicle values, *post hoc vs*. control, *p* = 0.0014) and MDG (GIRK1, n = 15 slices, 79 ± 4.7% of control vehicle values; *post hoc vs.* control, *p* < 0.001; GIRK2, n = 15, 81 ± 4.7% of control vehicle values, *post hoc vs*. control, *p* = 0.006). On the other hand, TQ induced a significant increase in GIRK1 and GIRK2 optical density (Fig. 4A, B, E) in both SLM (GIRK1, n = 16, 113 ± 3.8% of control vehicle values, *post hoc vs.* control vehicle, *p* = 0.0012; GIRK2, n = 15, 120 ± 5.9% of control vehicle values, *post hoc vs.* control, *p* = 0.0102) and MDG (GIRK1, n = 16 slices, 110 ± 5% of control values, *post hoc vs.* control vehicle, *p* = 0.044; GIRK2, n = 17, 113 ± 3.6% of control values, *post hoc vs*. control vehicle, *p* = 0.014). These results suggest that any modulation of basal GirK conductance may have a large impact in dorsal hippocampus channel protein expression.

**Figure 4.**
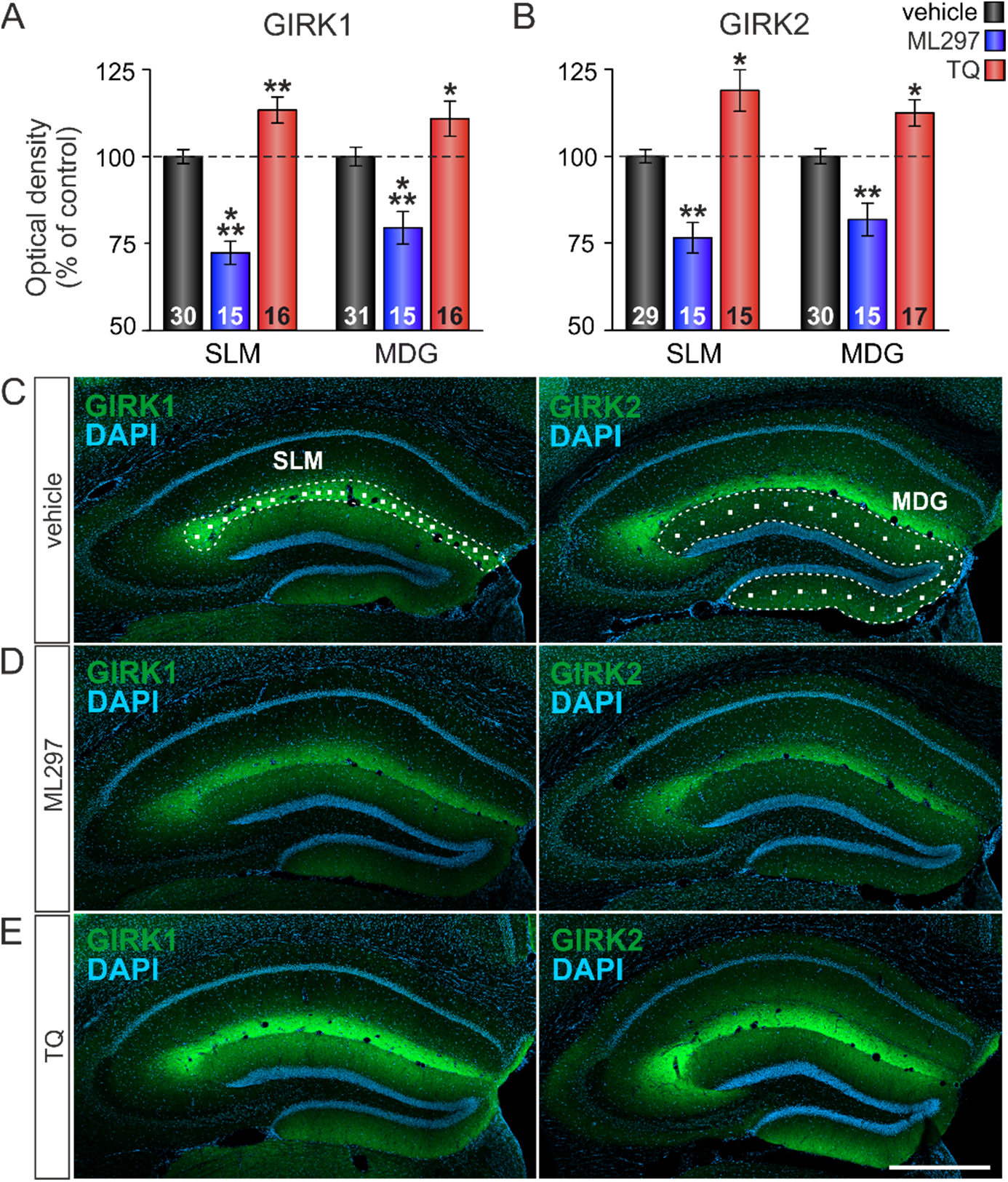
Hippocampal GirK protein expression pattern after basal GirK activity modulation. **(A, B)** Bar plots showing GIRK1 (A) and GIRK2 (B) immunostaining intensity (measured as optical density) expressed as percentage of control (vehicle) values (dashed line, 100%) in the *stratum lacunosum-moleculare* (SLM) and the molecular layer of the dentate gyrus (MDG) of the dorsal hippocampus in *i.c.v*. injected mice with vehicle (control), ML297 or TQ. Number of hippocampal sections for each condition is indicated in the corresponding bar (in animals: vehicle, n = 7; TQ, n = 4; ML297, n = 4). Differences *vs.* control (vehicle) are indicated by asterisks (*, *p* < 0.05; **, *p* < 0.01; ***, *p* < 0.001). (**C)**Confocal fluorescence photomicrographs showing the distribution of GIRK1 (left) and GIRK2 (right) subunits (green labelling) and DAPI stained cells (blue labelling) in the dorsal hippocampus of representative vehicle-injected mice. The images illustrate how random 15 × 15 μm squares were distributed through the SLM (left panel, area within the white dashed line) and MDG (right panel, area within the white dashed line) to measure GIRK1 and GIRK2 optical density. (**D, E)** Representative confocal microscopy images showing GIRK1 (left) and GIRK2 (right) immunostaining in *i.c.v.* injected mice with ML297 or TQ. Calibration bar: 500 μm.

Together, our results show, for the first time and at different levels of complexity, that basal GirK activity is required to perform cognitive functions that depends on dorsal hippocampus.

## Discussion

### Basal GirK activity, excitability, and synaptic plasticity at dorsal hippocampus

Our *ex vivo* and *in vivo* results showed an impaired LTP in the dorsal hippocampus when basal GirK activity is pharmacologically modulated. LTP was blocked when ML297 boosted GirK conductance very likely due to the high resting GirK conductance in the dorsal hippocampal dendrites, which decreases excitability and increases the threshold for LTP induction (Malik and Johnston, 2017). Moreover, the excess of hyperpolarization owing to GirK conductance enhancement may explain that HFS protocol led to insufficient activation of NMDA receptors, which results in a modest increase in intracellular Ca^2+^ levels, causing a decrease in synaptic efficacy (Cummings et al., 1996). In CA3–CA1 synapse of behaving mice, LTD has not been possible to evoke by classical 600 pulses presented at 1 Hz in slices (Bliss et al., 2006; Gruart et al., 2015). However, it can be induced by HFS if threshold required for LTP induction is not reached (for example GirK activity enhancement), according to the Bienenstock, Cooper and Munro (BCM) theory of synaptic plasticity that suggests the necessity of a certain threshold for LTP induction (Cooper and Bear, 2012). Hence, modulation of basal GirK activity may facilitate or compromise both plasticity processes, LTP/LTD. This is an important mechanism to produce changes in the ability to induce subsequent synaptic plasticity under physiological and pathological conditions, a basic phenomenon for metaplasticity (Abraham, 2008).

We also found that decreasing GirK conductance with TQ disrupts LTP *ex vivo* and *in vivo*. However, these results revealed significant differences with previous *ex vivo* studies where TQ produced an increase in LTP amplitude (Malik and Johnston, 2017). This discrepancy could be explained by several differences: *i*) in our *ex vivo* preparation, the depression of LTP in slices appeared at significant longer periods (>30 min) than the examined by them (first 30 min), and *ii*) we recorded *f*EPSPs from CA1 *stratum radiatum*, with greater sensitivity to detect changes in CA3–CA1 parameters (Gruart et al., 2012) than patch-clamping recordings obtained from one pyramidal cell. To assure these results, we performed a dose-response study and confirmed that LTP was always depressed by TQ. The explanation to this phenomenon might rely on a self-regulation mechanism of GABAergic interneurons that is crucial for the induction of LTP. During HFS, GABA reduces its own release by activating GABAB autoreceptors (*i.e.* generating a GirK activity increase), which assures enough NMDARs activation to induce LTP (Davies et al., 1991). Similarly, CA1 pyramidal neurons can release endocannabinoids that act as retrograde messengers to activate CB1 receptors, which are expressed primarily in GABAergic interneurons (Katona et al., 2000), also inhibiting GABA release (Wilson and Nicoll, 2001). As hippocampal GirK channels may act as effector of GABAB or CB1 receptors (Luscher and Slesinger, 2010), its blockade likely interferes with such regulation process, causing an excess of GABA in the medium, the inhibition on pyramidal neurons and LTP depression over time. The same situation might also be used to explain our *in vivo* results, as in behaving animals HFS intensity needs to be higher than *ex vivo* to successfully induce LTP (Madroñal et al., 2007), so in the presence of TQ, GABA release would be so high that NMDARs cannot properly be activated (Davies et al., 1991), also explaining the HFS–induced LTD (Keck et al., 2017).

The depressive effect of TQ on LTP *ex vivo* showed a slow time course noticeable after 30 min of potentiation, which suggests a differential role over the diverse temporal phases of synaptic plasticity. LTP can temporally and mechanistically be divided mainly into two phases, the early phase of LTP (E-LTP) and the late phase of LTP (L-LTP), functionally linked to short term memory (STM) and long term memory (LTM) processes (Bliss et al., 2007; Hardt et al., 2014). Our *ex vivo* data showed that although GirK activity was decreased by TQ, the synaptic response recorded from CA1 may be potentiated by HFS for at least 30 min. After this E–LTP phase, GirK activity deficits induced LTP–maintenance mechanisms to fail, and potentiation decayed. The conversion of E–LTP into L–LTP seems to be critical for the STM transformation into LTM (Villarreal et al., 2002). So the increased release of GABA discussed in the previous paragraph might induce a slow LTP suppression (Davies et al., 1991; Wilson and Nicoll, 2001) as it has recently been proposed for tonic GABA_A_ conductance in the hippocampus (Dembitskaya et al., 2020).

But in dorsal hippocampus GirK channels not only gate glutamatergic LTP (Malik and Johnston, 2017). LTP of GirK currents has been described (Huang et al., 2005) and recently proposed as a mechanism involved in the extinction of *f*EPSP potentiation to basal levels in order to processing new synaptic plasticity events and subsequent learning and memory formation, as *in vivo* it appeared 48h after HFS (Sanchez-Rodriguez et al., 2019). Interestingly, HFS elevates GirK channel surface density by NMDARs activation (Chung et al., 2009) explaining why such long-term inhibitory effects are more noticeable at longer times, when mechanisms needed to initiate the L–LTP maintenance take place (Hardt et al., 2014). In addition, synaptic plasticity can be modified by changing the level of pre- and postsynaptic GABA_B_ and A_1_ receptors desensitization via GirK activity (Wetherington and Lambert, 2002; Reis et al., 2019; Hill et al., 2020). These findings would be in agreement with our immunohistochemical findings, as sustained changes in the activity of the channel may or its trafficking regulation may induce compensatory regulations at the protein level.

Together, our data suggests that basal GirK activity may have a pivotal role in the control of threshold for LTP/LTD induction and plasticity maintenance mechanisms with significant impact on cognitive health.

### Basal GirK activity is essential for non-associative and associative dorsal hippocampus-dependent memories

When non-associative habituation learning was tested in the OF task, we found GirK modulation to produce habituation memory deficits in accordance with excitability and LTP impairments observed both *ex vivo* and *in vivo*, as animals kept exploring significantly more than controls. It has been recently shown that hippocampal hyperexcitability and plasticity deficits induced by acute amyloidopathy also impaired exploratory habituation in alert mice (Mayordomo-Cava et al., 2020; Sanchez-Rodriguez et al., 2020). Interestingly, metaplastic GirK activation reestablished LTP and hippocampal dependent memories (Sanchez-Rodriguez et al., 2017; Sanchez-Rodriguez et al., 2020) as also proposed in other amyloidosis models (Li et al., 2017; Peineau et al., 2018). Moreover, intraperitoneal administration of VU0466551, a 2-fold improved potency ML297 analogue, increased exploratory activity (Abney et al., 2019) while reducing GirK–dependent signaling constitutively (Pravetoni and Wickman, 2008), or selectively in dorsal hippocampus (Victoria et al., 2016), produced hyperactivity. Nevertheless, diffusion of *i.c.v*. injections have been shown to be mainly restricted to the hippocampal formation (Sanchez-Rodriguez et al., 2020) and *i.c.v.* administration of ML297 and TQ were proven not to alter locomotor activity (Sanchez-Rodriguez et al., 2020). Therefore, our results are not due to general motor effects of the drugs, and demonstrate the importance of hippocampal GirK conductance to sustain plasticity processes involved in exploratory habituation memory.

It has been reported that recognition memory and hippocampal synaptic plasticity processes are regulated through an inhibitory control mediated by 5-HT1A receptors (Fernandez et al., 2017). Such capability relies on correct CA3−CA1 synaptic functionality and subsequent LTP induction (Clarke et al., 2010). In fact, their selective pharmacological activation has also shown to facilitate depotentiation of synaptic strength (Fernandez et al., 2017), very likely through GirK channels in a similar mechanism as reported here. Furthermore, basal GirK conductance has been associated to A_1_R constitutive activity in the hippocampus (Reis et al., 2019; Hill et al., 2020). A_1_R activation induces a LTP depotentiation to prepare the hippocampus for subsequent synaptic processing and avoid synaptic overload (Izumi and Zorumski, 2019). It has been recently demonstrated *in vivo* that such mechanism requires of GirK channels to take place (Sanchez-Rodriguez et al., 2019). In fact, the inhibition of A_1_R has shown to block depotentiation in the CA1 region of the hippocampus, suggesting a role in memory erasure (Madroñal et al., 2016).

Finally, our results showed that modulation of basal GirK activity in the hippocampus also impacts on associative long-term memory as it was needed for developing a complex task as L/D operant test. The hippocampus is involved in both acquisition and storage of associative learning such as operant conditioning (Jurado-Parras et al., 2016). In fact, the lack of presynaptic GABAB receptors (of which GirK channels are main effectors) at hippocampal glutamatergic synapses, also impaired acquisition of an operant learning task (Jurado-Parras et al., 2016) suggesting the critical GirK channels contribution to certain forms of associative learning (Pravetoni and Wickman, 2008).

In summary, we found that basal GirK activity adjusts the dorsal hippocampus synapses to undertake the required changes for memory formation. When this metaplastic mechanism fails, marked deleterious changes are produced in hippocampal cognitive functions such as learning an associative instrumental task or recalling familiar objects and environments. In consequence, GirK channels gain relevance as one of the main determinants of neuronal excitability to support dorsal hippocampal-dependent cognitive functions.

## Acknowledgements

This work was supported by Spanish Ministry of Economy and Competitiveness from MINECO-FEDER (BFU2017-82494-P) to LJ-D and JDN-L, Fundación Tatiana Perez de Guzmán el Bueno to LJ-D, and “Plan Propio de Investigación” Programmes of University of Castilla-La Mancha (Predoctoral to IS-R and GI-L, and Senior Visiting Researchers to MON-M and AM). We thank Jose M. Gonzalez, María Sánchez and Jose A. Santos for their excellent technical assistance.

## Author Contributions

Conceived and designed the experiments: JDN-L, LJ-D. Performed the experiments: ST-C, SD, GI-L, IS-R, MONM and AM. Contributed materials: JMD-G and AG. Analyzed the data: ST-C, SD, GI-L, IS-R, JDN-L, and LJ-D. Wrote the paper: JDN-L and LJ-D. All authors revised the final version of the paper.

## Reference List

Abney KK, Bubser M, Du Y, Kozek KA, Bridges TM, Linsdley CW, Daniels JS, Morrison RD, Wickman K, Hopkins CR, Jones CK, Weaver CD (2019) Analgesic Effects of the GIRK Activator, VU0466551, Alone and in Combination with Morphine in Acute and Persistent Pain Models. ACS Chem Neurosci 10:1294–1299.

Abraham WC (2008) Metaplasticity: tuning synapses and networks for plasticity. Nat Rev Neurosci 9:387.

Bliss T, Collingridge GL, Morris R, Reymann KG (2018) Long-term potentiation in the hippocampus: discovery, mechanisms and function. Neuroforum 24:A103–A120.

Bliss TV, Gardner-Medwin AR (1973) Long-lasting potentiation of synaptic transmission in the dentate area of the unanaestetized rabbit following stimulation of the perforant path. J Physiol 232:357–374.

Bliss TV, Collingridge GL, Laroche S (2006) Neuroscience. ZAP and ZIP, a story to forget. Science 313:1058–1059.

Bliss TV, Collingridge GL, Morris R (2007) Synaptic Plasticity in the Hippocampus. In: The Hippocampus Book (Andersen P, Morris R, Amaral DG, Bliss T, O’Keefe J, eds), pp 343–474. New York: Oxford University Press.

Chen X, Johnston D (2005) Constitutively active G-protein-gated inwardly rectifying K+ channels in dendrites of hippocampal CA1 pyramidal neurons. J Neurosci 25:3787–3792.

Chung HJ, Ge WP, Qian X, Wiser O, Jan YN, Jan LY (2009) G protein-activated inwardly rectifying potassium channels mediate depotentiation of long-term potentiation. Proc Natl Acad Sci U S A 106:635–640.

Clarke JR, Cammarota M, Gruart A, Izquierdo I, Delgado-Garcia JM (2010) Plastic modifications induced by object recognition memory processing. Proc Natl Acad Sci USA 107:2652–2657.

Collingridge GL, Peineau S, Howland JG, Wang YT (2010) Long-term depression in the CNS. Nat Rev Neurosci 11:459–473.

Cooper LN, Bear MF (2012) The BCM theory of synapse modification at 30: interaction of theory with experiment. Nat Rev Neurosci 13:798–810.

Cummings JA, Mulkey RM, Nicoll RA, Malenka RC (1996) Ca2+ signaling requirements for long-term depression in the hippocampus. Neuron 16:825–833.

Dascal N, Kahanovitch U (2015) The Roles of Gbetagamma and Galpha in Gating and Regulation of GIRK Channels. Int Rev Neurobiol 123:27–85.

Davies CH, Starkey SJ, Pozza MF, Collingridge GL (1991) GABA autoreceptors regulate the induction of LTP. Nature 349:609–611.

Dembitskaya Y, Wu YW, Semyanov A (2020) Tonic GABAA Conductance Favors Spike-Timing-Dependent over Theta-Burst-Induced Long-Term Potentiation in the Hippocampus. J Neurosci 40:4266–4276.

Drake CT, Bausch SB, Milner TA, Chavkin C (1997) GIRK1 immunoreactivity is present predominantly in dendrites, dendritic spines, and somata in the CA1 region of the hippocampus. Proc Natl Acad Sci USA 94:1007–1012.

Fanselow MS, Dong HW (2010) Are the dorsal and ventral hippocampus functionally distinct structures? Neuron 65:7–19.

Fernandez-Alacid L, Watanabe M, Molnar E, Wickman K, Lujan R (2011) Developmental regulation of G protein-gated inwardly-rectifying K+ (GIRK/Kir3) channel subunits in the brain. Eur J Neurosci 34:1724–1736.

Fernandez-Lamo I, Delgado-Garcia JM, Gruart A (2018) When and Where Learning is Taking Place: Multisynaptic Changes in Strength During Different Behaviors Related to the Acquisition of an Operant Conditioning Task by Behaving Rats. Cereb Cortex 28:1011–1023.

Fernandez SP, Muzerelle A, Scotto-Lomassese S, Barik J, Gruart A, Delgado-Garcia JM, Gaspar P (2017) Constitutive and Acquired Serotonin Deficiency Alters Memory and Hippocampal Synaptic Plasticity. Neuropsychopharmacology 42:512–523.

Glaaser IW, Slesinger PA (2015) Structural Insights into GIRK Channel Function. Int Rev Neurobiol 123:117–160.

Gonzalez C, Baez-Nieto D, Valencia I, Oyarzun I, Rojas P, Naranjo D, Latorre R (2012) K(+) channels: function-structural overview. Compr Physiol 2:2087–2149.

Gruart A, Delgado-Garcia JM (2007) Activity-dependent changes of the hippocampal CA3-CA1 synapse during the acquisition of associative learning in conscious mice. Genes Brain Behav 6 Suppl 1:24–31.

Gruart A, Munoz MD, Delgado-Garcia JM (2006) Involvement of the CA3-CA1 synapse in the acquisition of associative learning in behaving mice. J Neurosci 26:1077–1087.

Gruart A, Benito E, Delgado-Garcia JM, Barco A (2012) Enhanced cAMP response element-binding protein activity increases neuronal excitability, hippocampal long-term potentiation, and classical eyeblink conditioning in alert behaving mice. J Neurosci 32:17431–17441.

Gruart A, Leal-Campanario R, Lopez-Ramos JC, Delgado-Garcia JM (2015) Functional basis of associative learning and its relationships with long-term potentiation evoked in the involved neural circuits: Lessons from studies in behaving mammals. Neurobiol Learn Mem 124:3–18.

Hardt O, Nader K, Wang YT (2014) GluA2-dependent AMPA receptor endocytosis and the decay of early and late long-term potentiation: possible mechanisms for forgetting of short- and long-term memories. Philos Trans R Soc Lond B Biol Sci 369:20130141.

Hasan MT, Hernandez-Gonzalez S, Dogbevia G, Trevino M, Bertocchi I, Gruart A, Delgado-Garcia JM (2013) Role of motor cortex NMDA receptors in learning-dependent synaptic plasticity of behaving mice. Nat Commun 4:2258.

Hill E, Hickman C, Diez R, Wall M (2020) Role of A1 receptor-activated GIRK channels in the suppression of hippocampal seizure activity. Neuropharmacology 164:107904.

Huang CS, Shi SH, Ule J, Ruggiu M, Barker LA, Darnell RB, Jan YN, Jan LY (2005) Common molecular pathways mediate long-term potentiation of synaptic excitation and slow synaptic inhibition. Cell 123:105–118.

Izumi Y, Zorumski CF (2019) Temperoammonic Stimulation Depotentiates Schaffer Collateral LTP via p38 MAPK Downstream of Adenosine A1 Receptors. J Neurosci 39:1783–1792.

Jurado-Parras MT, Delgado-Garcia JM, Sanchez-Campusano R, Gassmann M, Bettler B, Gruart A (2016) Presynaptic GABAB Receptors Regulate Hippocampal Synapses during Associative Learning in Behaving Mice. PLoS One 11:e0148800.

Katona I, Sperlagh B, Magloczky Z, Santha E, Kofalvi A, Czirjak S, Mackie K, Vizi ES, Freund TF (2000) GABAergic interneurons are the targets of cannabinoid actions in the human hippocampus. Neuroscience 100:797–804.

Keck T, Hubener M, Bonhoeffer T (2017) Interactions between synaptic homeostatic mechanisms: an attempt to reconcile BCM theory, synaptic scaling, and changing excitation/inhibition balance. Curr Opin Neurobiol 43:87–93.

Leussis MP, Bolivar VJ (2006) Habituation in rodents: a review of behavior, neurobiology, and genetics. Neurosci Biobehav Rev 30:1045–1064.

Li Q, Navakkode S, Rothkegel M, Soong TW, Sajikumar S, Korte M (2017) Metaplasticity mechanisms restore plasticity and associativity in an animal model of Alzheimer’s disease. Proc Natl Acad Sci U S A 114:5527–5532.

Lujan R, Aguado C (2015) Localization and Targeting of GIRK Channels in Mammalian Central Neurons. Int Rev Neurobiol 123:161–200.

Lujan R, Marron Fernandez de Velasco E, Aguado C, Wickman K (2014) New insights into the therapeutic potential of Girk channels. Trends Neurosci 37:20–29.

Luscher C, Slesinger PA (2010) Emerging roles for G protein-gated inwardly rectifying potassium (GIRK) channels in health and disease. Nat Rev Neurosci 11:301–315.

Luscher C, Malenka RC (2012) NMDA receptor-dependent long-term potentiation and long-term depression (LTP/LTD). Cold Spring Harb Perspect Biol 4.

Madroñal N, Delgado-Garcia JM, Gruart A (2007) Differential effects of long-term potentiation evoked at the CA3 CA1 synapse before, during, and after the acquisition of classical eyeblink conditioning in behaving mice. J Neurosci 27:12139–12146.

Madroñal N, Delgado-Garcia JM, Fernandez-Guizan A, Chatterjee J, Kohn M, Mattucci C, Jain A, Tsetsenis T, Illarionova A, Grinevich V, Gross CT, Gruart A (2016) Rapid erasure of hippocampal memory following inhibition of dentate gyrus granule cells. Nat Commun 7:10923.

Malik R, Johnston D (2017) Dendritic GIRK Channels Gate the Integration Window, Plateau Potentials, and Induction of Synaptic Plasticity in Dorsal But Not Ventral CA1 Neurons. J Neurosci 37:3940–3955.

Malleret G, Alarcon JM, Martel G, Takizawa S, Vronskaya S, Yin D, Chen IZ, Kandel ER, Shumyatsky GP (2010) Bidirectional regulation of hippocampal long-term synaptic plasticity and its influence on opposing forms of memory. J Neurosci 30:3813–3825.

Mayfield J, Blednov YA, Harris RA (2015) Behavioral and Genetic Evidence for GIRK Channels in the CNS: Role in Physiology, Pathophysiology, and Drug Addiction. Int Rev Neurobiol 123:279–313.

Mayordomo-Cava J, Iborra-Lazaro G, Djebari S, Temprano-Carazo S, Sanchez-Rodriguez I, Jeremic D, Gruart A, Delgado-Garcia JM, Jimenez-Diaz L, Navarro-Lopez JD (2020) Impairments of Synaptic Plasticity Induction Threshold and Network Oscillatory Activity in the Hippocampus Underlie Memory Deficits in a Non-Transgenic Mouse Model of Amyloidosis. Biology (Basel) 9.

Morris R (2007) Theories of Hippocampal Function. In: The Hippocampus Book (Andersen P, Morris R, Amaral DG, Bliss T, O’Keefe J, eds), pp 581–713. New York: Oxford University Press.

Nava-Mesa MO, Jimenez-Diaz L, Yajeya J, Navarro-Lopez JD (2013) Amyloid-beta induces synaptic dysfunction through G protein-gated inwardly rectifying potassium channels in the fimbria-CA3 hippocampal synapse. Front Cell Neurosci 7:117.

Nava-Mesa MO, Jimenez-Diaz L, Yajeya J, Navarro-Lopez JD (2014) GABAergic neurotransmission and new strategies of neuromodulation to compensate synaptic dysfunction in early stages of Alzheimer’s disease. Front Cell Neurosci 8:167.

Paxinos G, Franklin KB (2001) The Mouse Brain in Stereotaxic Coordinates. London: Academic Press.

Peineau S, Rabiant K, Pierrefiche O, Potier B (2018) Synaptic plasticity modulation by circulating peptides and metaplasticity: Involvement in Alzheimer’s disease. Pharmacol Res 130:385–401.

Pravetoni M, Wickman K (2008) Behavioral characterization of mice lacking GIRK/Kir3 channel subunits. Genes Brain Behav 7:523–531.

Reis SL, Silva HB, Almeida M, Cunha RA, Simoes AP, Canas PM (2019) Adenosine A1 and A2A receptors differently control synaptic plasticity in the mouse dorsal and ventral hippocampus. J Neurochem.

Rifkin RA, Moss SJ, Slesinger PA (2017) G Protein-Gated Potassium Channels: A Link to Drug Addiction. Trends Pharmacol Sci 38:378–392.

Sanchez-Rodriguez I, Gruart A, Delgado-Garcia JM, Jimenez-Diaz L, Navarro-Lopez JD (2019) Role of GirK Channels in Long-Term Potentiation of Synaptic Inhibition in an In Vivo Mouse Model of Early Amyloid-beta Pathology. Int J Mol Sci 20.

Sanchez-Rodriguez I, Temprano-Carazo S, Najera A, Djebari S, Yajeya J, Gruart A, Delgado-Garcia JM, Jimenez-Diaz L, Navarro-Lopez JD (2017) Activation of G-protein-gated inwardly rectifying potassium (Kir3/GirK) channels rescues hippocampal functions in a mouse model of early amyloid-beta pathology. Sci Rep 7:14658.

Sanchez-Rodriguez I, Djebari S, Temprano-Carazo S, Vega-Avelaira D, Jimenez-Herrera R, Iborra-Lazaro G, Yajeya J, Jimenez-Diaz L, Navarro-Lopez JD (2020) Hippocampal long-term synaptic depression and memory deficits induced in early amyloidopathy are prevented by enhancing G-protein-gated inwardly rectifying potassium channel activity. J Neurochem 153:362–376.

Slesinger, P.A., Wickman, K. (2015) Structure to Function of G Protein-Gated Inwardly Rectifying (GIRK) Channels, 1^a^ Edition: Elsevier Inc.

Tipps ME, Buck KJ (2015) GIRK Channels: A Potential Link Between Learning and Addiction. Int Rev Neurobiol 123:239–277.

Trompoukis G, Rigas P, Leontiadis LJ, Papatheodoropoulos C (2020) Ih, GIRK, and KCNQ/Kv7 channels differently modulate sharp wave - ripples in the dorsal and ventral hippocampus. Mol Cell Neurosci 107:103531.

Victoria NC, Marron Fernandez de Velasco E, Ostrovskaya O, Metzger S, Xia Z, Kotecki L, Benneyworth MA, Zink AN, Martemyanov KA, Wickman K (2016) G Protein-Gated K+ Channel Ablation in Forebrain Pyramidal Neurons Selectively Impairs Fear Learning. Biol Psychiatry 80:796–806.

Villarreal DM, Do V, Haddad E, Derrick BE (2002) NMDA receptor antagonists sustain LTP and spatial memory: active processes mediate LTP decay. Nat Neurosci 5:48–52.

Wetherington JP, Lambert NA (2002) Differential desensitization of responses mediated by presynaptic and postsynaptic A1 adenosine receptors. J Neurosci 22:1248–1255.

Whorton MR, MacKinnon R (2013) X-ray structure of the mammalian GIRK2-betagamma G-protein complex. Nature 498:190–197.

Wilson RI, Nicoll RA (2001) Endogenous cannabinoids mediate retrograde signalling at hippocampal synapses. Nature 410:588–592.

Xiong G, Metheny H, Johnson BN, Cohen AS (2017) A Comparison of Different Slicing Planes in Preservation of Major Hippocampal Pathway Fibers in the Mouse. Front Neuroanat 11:107.

Yow TT, Pera E, Absalom N, Heblinski M, Johnston GA, Hanrahan JR, Chebib M (2011) Naringin directly activates inwardly rectifying potassium channels at an overlapping binding site to tertiapin-Q. BrJPharmacol 163:1017–1033.

Zucker RS, Regehr WG (2002) Short-term synaptic plasticity. Annu Rev Physiol 64:355–405.

